# TLR4/NF-κB Signaling Contributes to the Inflammation in Ovary and Inguinal Fats of Polycystic Ovary Syndrome

**DOI:** 10.1101/2022.07.14.500077

**Authors:** Bo Sun, Zimeng Pan, Hongying Kuang, Xiaoling Feng, Lihui Hou, Fang Xu, Feifei Song, Miao Sun

## Abstract

**Background:** Polycystic ovary syndrome (PCOS) is a complex reproductive endocrine disorder. It has highly heterogeneous clinical manifestations which are characterized by biochemical hyperandrogenemia, obesity, insulin resistance, dyslipidemia, anovulation, and polycystic ovaries. However, the etiologies of PCOS are still unclear. Recently, studies have found that the low-grade inflammation contributed to the occurrence of PCOS, and as a critical biomarker indicated the endocrine disruptions in PCOS.

**Objective:** This study is aimed to investigate the processes and mediators of inflammation in contributing to the development of PCOS.

**Methods:** Letrozole (LET) induced PCOS rat model was used in this study. Body weight, body temperature, inguinal fats weight, fasting glucose level, ovarian morphology, NF-κB signaling target genes in ovary, and protein expression levels of TLR4 and NF-κB in ovarian and inguinal fats were measured in rats with placebo and LET administrations for 6 and 12 weeks.

**Results:** PCOS rats, especially with LET intervention for 12 weeks, had higher body weight, inguinal fats weight and fasting glucose level compared to control group. The protein expression levels of TLR4 and NF-κB in cytoplasm of ovarian and inguinal fats were increased in LET-induced PCOS rats compared to control groups, while NF-κB in nucleus were reduced in PCOS rats. The expressions of ACTB, C3, CXCL3, NQO1 and SELP in ovarian were statistically different in PCOS rats induced by LET compared to control groups.

**Conclusion:** These findings indicated that stimulating TLR4/NF-κB pathway in inguinal fats and ovary tissues contributed to the increased inflammation in LET-induced PCOS rats, which, in turn, exacerbated the phenotype of PCOS including weight gain, adipose tissue accumulation, hyperglycemia and follicular dysplasia.

## 1 Introduction

Polycystic ovary syndrome (PCOS) is the most common endocrine disorder which affects 4%-20% women of reproductive age in the worldwide [1]. It is characterized by biochemical and/or clinical hyperandrogenemia, chronic anovulation, and polycystic ovaries [2]. PCOS usually occurred with other accompany symptoms including insulin resistance, dyslipidemia and abdominal obesity. It contributes to increased risks of developing metabolic syndrome which is a disease covering type 2 diabetes, cardiovascular diseases and hypertension [3]. PCOS severely influence the people’s health and brought essential social and economic burdens. However, the underlying mechanism is still unclear.

Studies have found PCOS patients have higher levels of inflammatory markers (C-reactive protein, interleukin 6 [IL-6], interleukin 18 and tumor necrosis factor-α [TNF-α]) compared with age- and BMI-matched healthy controls [4–6]. Studies suggest that chronic low-grade inflammation is associated with metabolic disorder and ovarian dysfunction in women with PCOS [7–9]. The chronic low-grade inflammation in PCOS women is found to be associated with hypertrophy adipocyte in which the compressed stromal vessels in adipose tissue causes poor blood supply leading to adipose tissue hypoperfusion and, consequently, hypoxia. Then cytokines were secreted and released in local adipose tissue [10]. Interestingly, the insulin resistance, dyslipidemia, and hyperandrogenism might be also in connection with lipid-induced proinflammatory response in PCOS [11]; study also suggests that the adipose tissue as a potential producer of these cytokines (e.g., TNF-α, leptin and resistin) contributing to this inflammation in PCOS [10]. Further, visceral adipose tissue contribute to maintaining this inflammation through regulating the secretions of cytokines and monocyte chemoattractant proteins, as well as recruiting more immune cells [9]. Therefore, chronic inflammation has been increasingly recognized as a critical target in studying the etiologies of PCOS.

PCOS can severely affect the follicle quality and oocyte maturation, in which cause infertility in women [12]. Inflammation plays an adverse effect on ovulation which are involved in premature ovarian failure, physiological aging of the ovaries and PCOS [9]. It found that the local inflammatory microenvironment in the ovary lead to abnormal follicular development and ovulation disorders [13]. It is also known that the inflammation in circulation plays an adverse effect on oocytes through altering follicle microenvironment, oxidative stress and granulocyte macrophage colony stimulating factors [14]. Therefore, the development of local inflammatory in the ovary as a key in studying PCOS needs more attentions.

Up to now, animal PCOS models have been developed efficiently with the characteristic features of PCOS; the most extensively reported is using testosterone, letrozole (LET) and estradiol valerate induced rats and mice model [15]. An animal model which can exhibit all symptoms of PCOS is the most desirable, since PCOS is a heterogeneous disease. LET is a nonsteroidal aromatase inhibitor [16], and can be administrated through oral or subcutaneous [17]. Our previous study indicated that rats with administration of LET subcutaneously for 12 weeks showed both the reproductive and metabolic alterations as PCOS patients have [18]. Although many studies have used LET-induced animal models to study the pathogenesis and pharmacological treatment related to PCOS, most of them focus on the characteristics related to the end induction of the model [19–24], and only a few experiments have been conducted to investigate the pathological mechanisms of LET-induced PCOS model animals at different time points [25,26]. Further, few studies work on investigating the regulation of inflammatory response during the pathogenesis of PCOS.

Therefore, this study intends to investigate the inflammatory processes and mediators that contribute to the commencement and development of PCOS across two different timepoints. By doing so it can broaden our understanding in the association between PCOS and inflammation, provide new direction on preventing and delaying the process of PCOS.

## 2 Materials and Methods

### Animals

Female Wistar rats were purchased from Experimental Animal Center of University of Heilongjiang Chinese Medicine (Harbin, China). All experimental procedures involving rats were approved by the University of Heilongjiang Chinese Medicine Animal Care and Use Committee and conformed to the Animal Welfare Act 1999. Briefly, female Wistar rats were housed under controlled conditions (21-22°C and 12 h light, 12 h dark cycle). Rats were fed on commercial chow and tap water ad libitum. On age of 21 days, female Wistar rats were randomly assigned to four groups: LET group 1 (LET1) (N=3, 200 μg/day; 60-day continuous-release LET pellets), LET group 2 (LET2) (N=3, 200 μg/day; 90-day continuous-release LET pellets), control group 1 (Con1) (N=3, 200 μg/day; 60-day continuous-release placebo pellets, the main ingredient of which is starch), and control group 2 (Con2) (N=3, 200 μg/day; 90-day continuous-release placebo pellets). LET (HY-14248, MedChemExpress, USA) and placebo (Innovative Research of America, USA) were implanted with subcutaneous either 60 days or 90 days continuous-release pallets (Innovative Research of America, Sarasota, FL). The 200 μg /day dose was chosen based on previous (18). The flow chart of this study is shown in Figure 1. Food intake, body weight and body temperature were measured every day and calculated weekly from the second week of implantation. Rats were culled and tissues were collected at 6 weeks of LET1 and Con1 and 12 weeks of LET2 and Con12 after pellet implantation, respectively.

**Figure 1.**
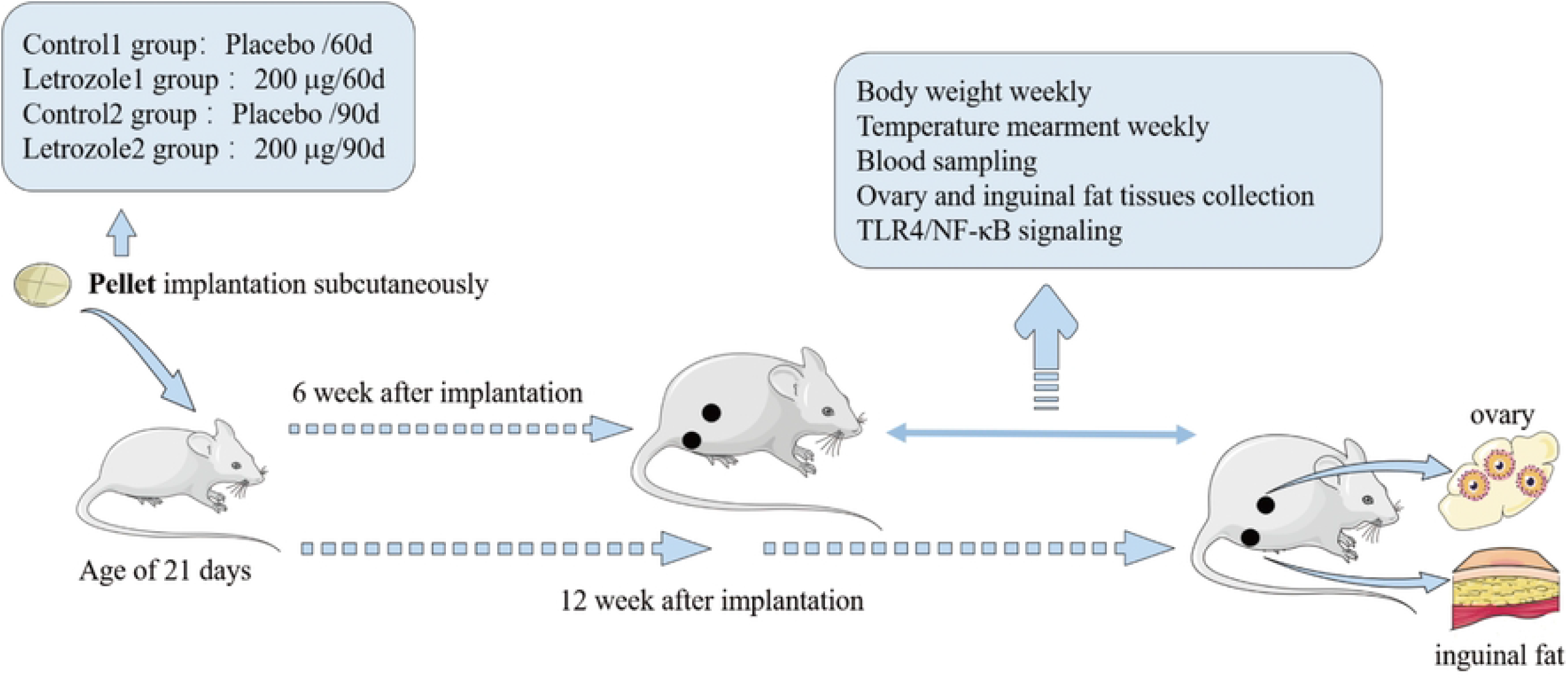

### Fasting Blood Glucose Assays

Rats were fasted overnight, and 1 ml of blood was collected from the tail vein. Blood samples were centrifuged (3000 rpm, 10 min, 4°C) (Remi, C-24 BL, Mumbai, India). Glucose in serum was measured by Glucose Oxidase and Peroxidase (GOD-POD) kit according to the manufacture’s instruction.

### H&E staining and Immunohistochemistry

Rats were culled by isoflurane asphyxiation for tissues collection after 6 and 12 weeks of LET administration. Ovaries and inguinal fats were collected freshly and weighed. One ovary and 0.5 cm length segment of inguinal fats were used for histology and the other ovary and last inguinal fats were stored at -80°C until processing for Western Blot. To observe ovary and fat morphology changes, hematoxylin and eosin (H&E) staining was used. Ovary and inguinal fat pads were fixed in 10% neutral buffered formalin for 24h and stored in 70% ethanol before performing H&E staining and immunohistochemistry. For H&E staining, tissues were embedded in paraffine using standard techniques.

Ovaries and inguinal fat pads were respectively, serially sectioned at 5 μm and then stained with hematoxylin and eosin. Images were collected using Olympus DP73 microscope. For immunohistochemistry, the protocol was described according to Shaaban’
ss study (27). Briefly, the ovary and inguinal fat sections were de-waxed, rehydrated, and performed antigen retrieval in 0.1 M sodium citrate buffer for 10 min in a microwave. Sections were incubated in 3% hydrogen peroxide for 15 min to eliminate endogenous peroxidases. Sections were blocked with goat serum for 15 min at 37 °C followed by incubation overnight at 4 °C with primary antibodies Toll-like receptor 4 (TLR4) (1: 200, WL00196, wanleibio, China) and nuclear factor-κB (NF-κB) (1: 200, WL01980, wanleibio, China). Following three washes in PBS, sections were incubated with biotin-labeled goat anti-rabbit IgG (H+L) (1: 500, #31460, ThermoFisher, USA) for 30 min at 37 °C. Signals were visualized by reaction with the DAB peroxidase substrate kit (DA1010, Solarbio, China). Images were collected using Olympus DP73 microscope.

### Western Bolt

30 mg ovaries and inguinal fats were respectively used for protein extraction. 40 µg total protein per sample was loaded onto the gel. The methods and procedures were described in previous [28]. The primary antibody we used are TLR4 (1:1000, WL00196; Wanleibio, China) and NF-κB P65 (1:500, WL01980; Wanleibio, China). Goat anti-rabbit IgG-horseradish peroxidase (HRP) (1:5,000, WLA023; Wanleibio, China) was used as secondary antibody. β-actin was used to normalize the expressions of either TLR4 or NF-κB in each sample. The intensities of the protein signals were detected and quantified by Gel-Pro-Analyzer (WD-9413B; Liuyi Biology, China).

### qRT-PCR Array Analysis

92 related genes expression profiles were measured using rat NF-κB Signaling Targets qPCR Array (Supplementary TableS1) according to the manufacturer’s protocol (Wcgene Biotech, Shanghai, China). Data was validated using real time quantitative reverse transcription PCR (qRT-PCR). Total RNA (1 μg) was extracted using Trizol and proceed to reverse transcription reactions using the High Capacity cDNA Reverse Transcription kit (Applied Biosystems, Foster City, CA). qRT-PCR was performed on a 7900HT Sequence Detection system (Applied Biosystems) using FastStart Universal SYBR Green Master kit (Roche Diagnostics, Indianapolis, IN) following the manufacturer’s instructions. GAPDH was used as a housekeeping gene. Each test was done in triple replication and the 2^−ΔΔCt^ method was used to calculate the expressions of genes. Data were analyzed using Wcgene Biotech software. Genes with fold-changes more than or less than 2.0 were considered to be of biological significance.

### Statistical Analysis

Social sciences (SPSS version 21.0; SPSS, Chicago, IL) and Prism GraphPad (version 6.00, GraphPad Software, La Jolla, CA) were used for testing the statistical difference. Differences between controls and rats exposed to LET were assessed by one-way ANOVA after Dunnett’s post hoc test. The differences of body weight, body temperature and food intake between groups were analyzed by repeated-measures ANOVA. Values are mean ± S.E.M. *P* < 0.05 was defined as the threshold of significant.

## 3 Results

### Body Weight Accumulation and Temperature Changes

The continuously released pellet containing placebo or LET were implanted subcutaneously at age of 21 day. The accumulation of body weight was recorded for 6 weeks and 12 weeks as shown in Figure 2A. Significantly increased body weight was observed in rats at 5 and 6 weeks after LET implantation (Con1 vs LET1, *P*<0.05). In addition, the body weight of LET2 group were increased compared with the Con 2 group during 7-12 weeks, *P*<0.05. Further, the changes of body temperature were fluctuated in the Con1 and Con2 groups, while the body temperature of LET2 rats were lower than Con2 group at the 7th, 8th and 9th week (Con2 vs LET2, *P*<0.05), as shown in Figure 2B.

**Figure 2.**
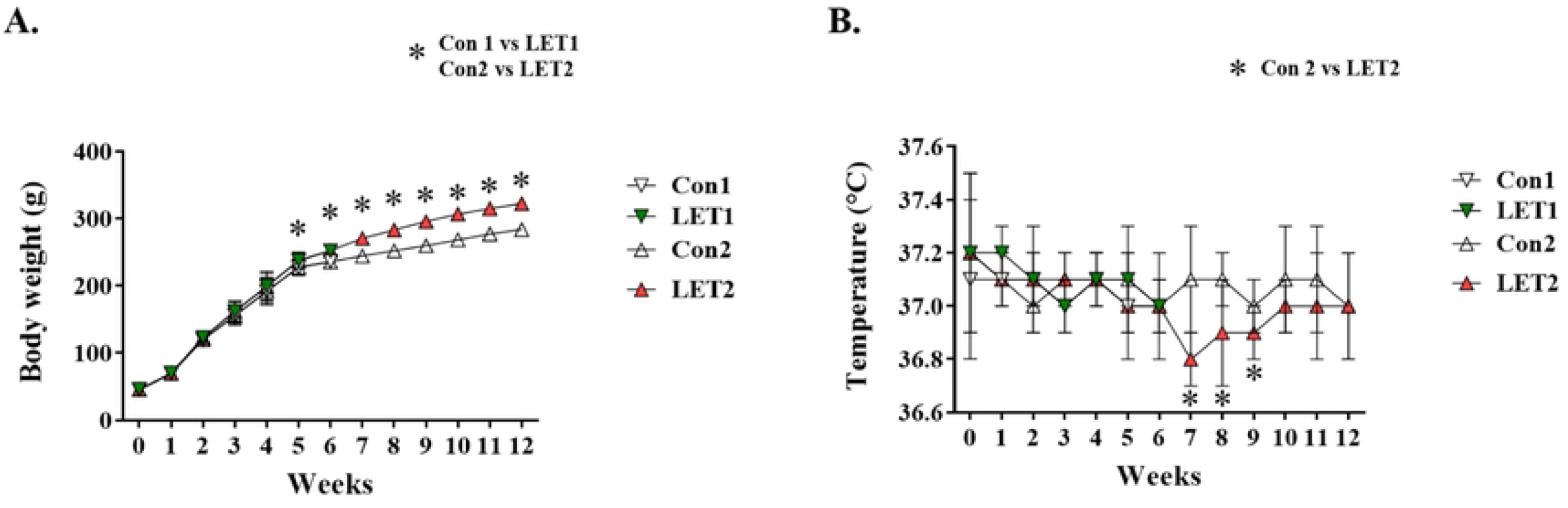

### Inguinal Fats Weight and Fasting Blood Glucose Level

The weights of inguinal fats tissue in four groups were compared and shown in Figure 3A. The weight of inguinal fats was increased as the treatment going (Con1 vs Con2, *P*<0.05). In addition, the weight of inguinal fats in rats at 6 and 12 weeks of LET implantation was higher than control group at the same period (Con1 vs LET1, *P* < 0.05; Con2 vs LET2, *P* < 0.001), and the inguinal fats weight at 12 weeks of LET implantation was higher than that at 6 weeks (*P* < 0.001). This suggested that the pathological state of PCOS was associated with the increased inguinal fats tissue weight. As the severity of PCOS increased, inguinal fats tissue also gradually got weight.

Figure 3B showed the fasting blood glucose level of the four groups. The fasting blood glucose level of rats at 6 and 12 weeks of LET implantation was higher than their corresponding control groups (Con1 vs LET1, *P* < 0.001; Con2 vs LET2, *P* < 0.001). And the fasting blood glucose level at 12 weeks of LET implantation was higher than that at 6 weeks (*P* < 0.01). This suggested that the risk of having glucose metabolism disorders was significantly increased in PCOS and became more severe as PCOS disease progresses.

**Figure 3.**
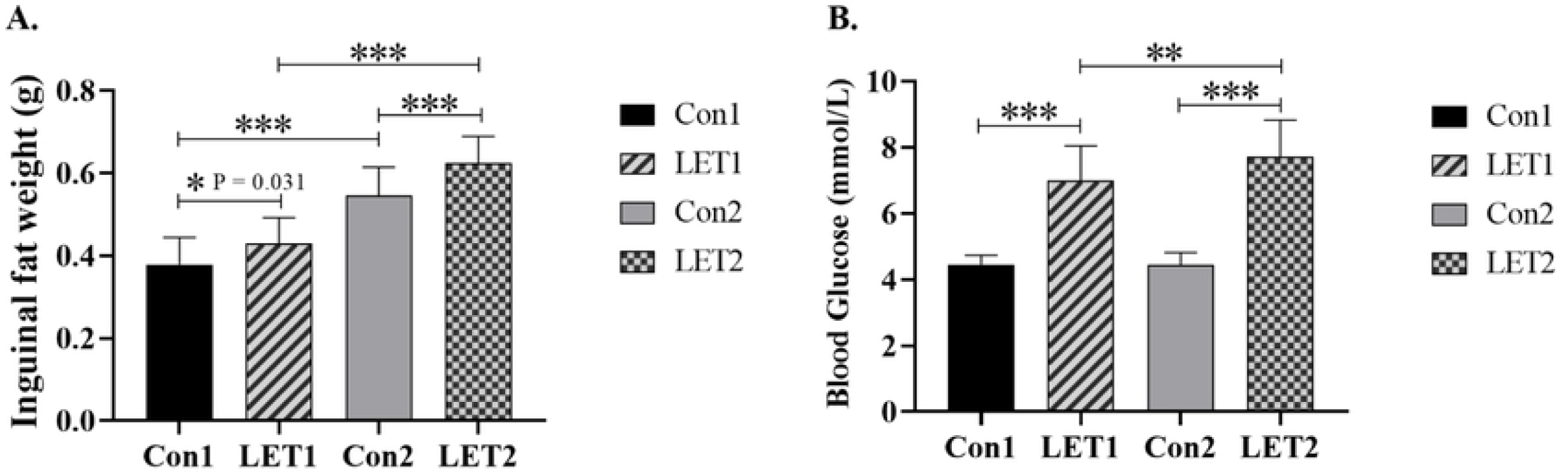

### Ovarian Morphology Changes

More follicles were observed in the ovarian cortex of Con1, Con2 and LET1 groups compared with LET2 group (Figure 4A, B and C). Luteum was clear, cumulus and oocytes were visible, granulosa cell layers were mostly 8 ∼ 9 layers and closely arranged, and mature follicles were intact without hyperemia and edema in Con1, Con2 and LET1 groups. The ovary of LET2 had cystic changes, with obvious cystic dilatation of follicles, more atresia, and reduced number of granulosa cell layers and luteal body (Figure 4 D).

**Figure 4.**
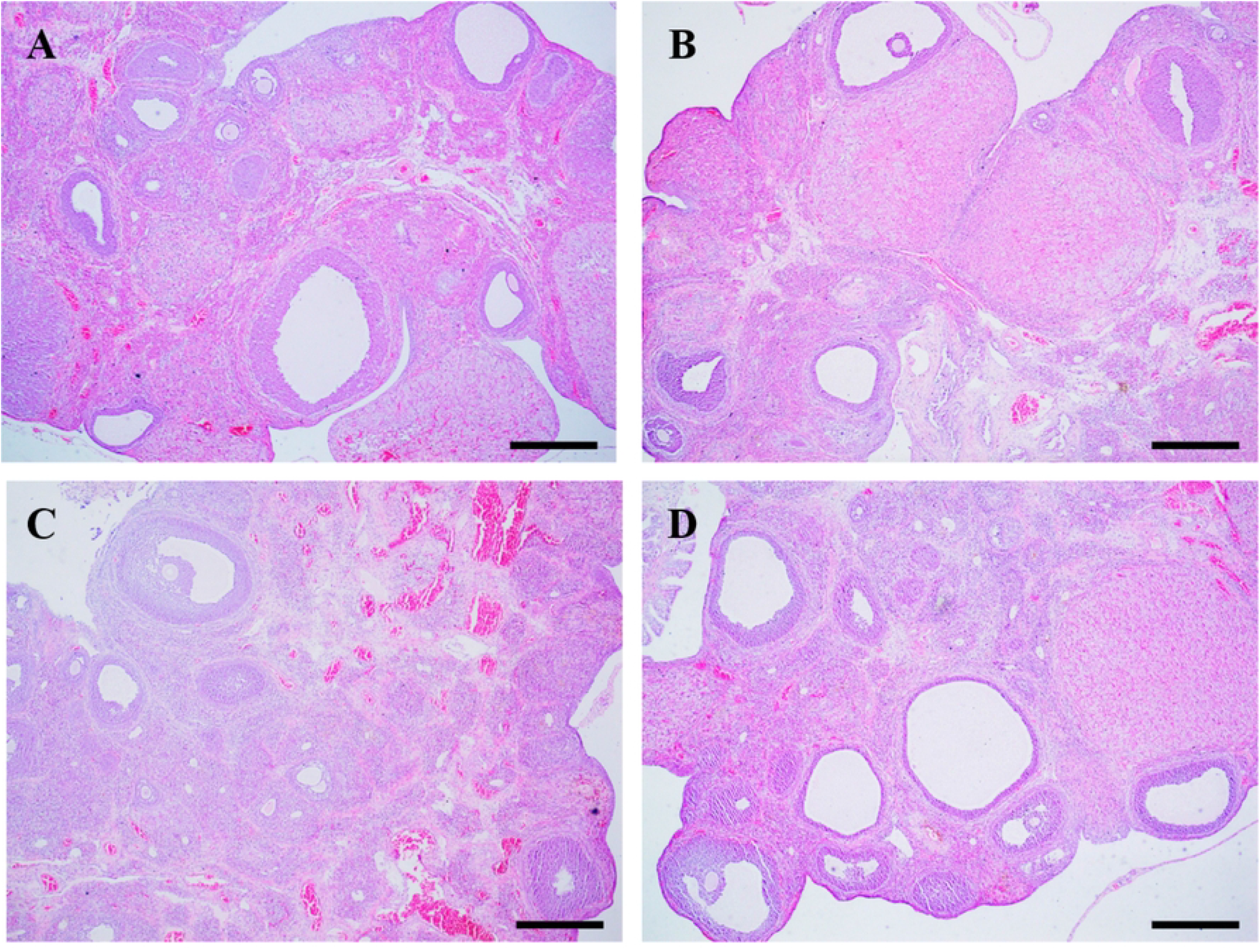

### The TLR4 and NF-κB Protein Expression in Ovaries and Inguinal Fats

The abundance of TLR4 and NF-κB protein expressions in ovaries and inguinal fat tissues were measured using immunohistochemistry and western blot. We found that the intensity of TLR4 and NF-κB immunostaining were increased in both ovaries (Figure 5D and Figure 6D) and inguinal fat tissues (Figure 5H and Figure 6H) in LET2. Consistently, western blot showed that the relative expression of TLR4 protein in ovaries and inguinal fat tissues were increased in LET groups compared to control groups (Con1 vs LET1, *P* < 0.05; Con2 vs LET2, *P* < 0.05, Figure 5I). And an increased expression of TLR4 was observed in LET2 compared with LET1 (LET2 vs LET1, *P* < 0.05, Figure 5I). Moreover, the protein expression of NF-κB in cytoplasm of ovaries and inguinal fat tissues were also increased in LET groups compared to control groups (Con1 vs LET1, *P* < 0.05; Con2 vs LET2, *P* < 0.05, Figure 6I). A greater increase in NF-κB expression in cytoplasm was also seen in LET2 compared to LET1 group (LET2 vs LET1, *P* < 0.05, Figure 6I). While the expression of NF-κB in nucleus was reduced in LET groups compared to their control groups in ovaries and inguinal tissues (Con1 vs LET1, *P* < 0.05; Con2 vs LET2, *P* < 0.05, Figure 6J). What’
ss more, there was a greater decrease in NF-κB expression in nucleus in LET2 compared to LET1 group (LET2 vs LET1, *P* < 0.05, Figure 6J).

**Figure 5.**
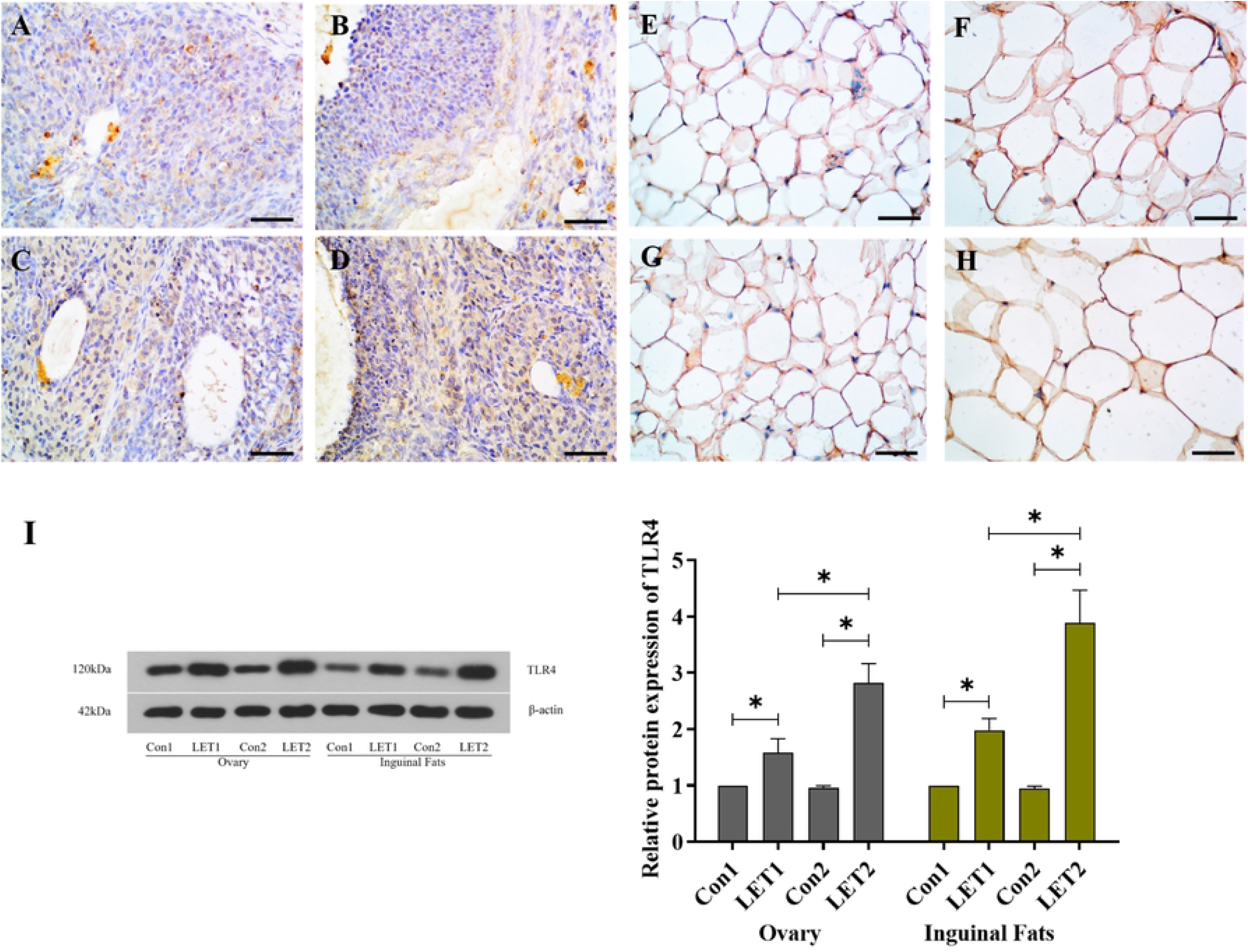

**Figure 6.**
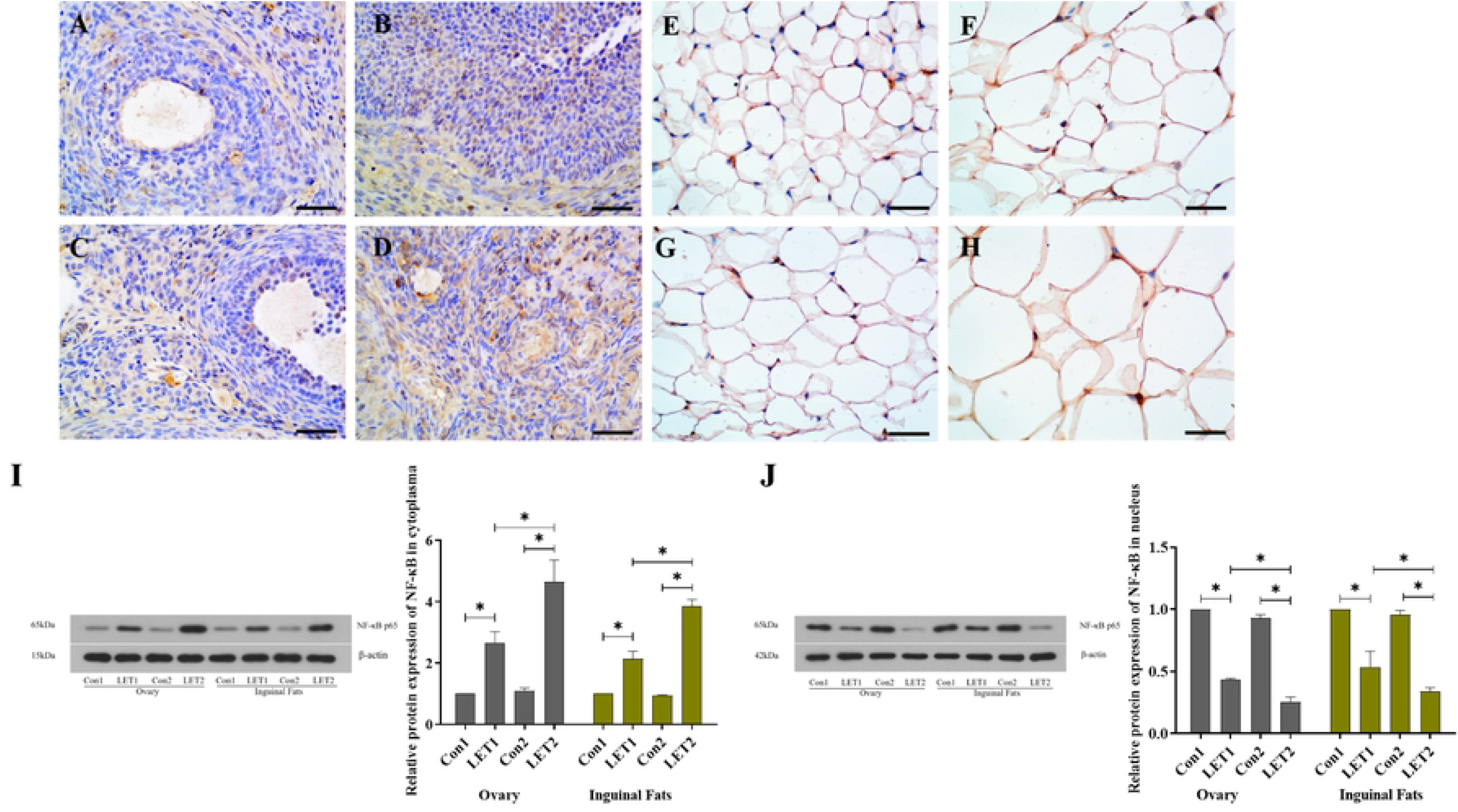

### The PCR array of NF-κB signaling targets in ovary

The genes involved in the NF-κB signaling were investigated using qRT-PCR and the results were shown in Supplementary data (TableS2). There were significant differences in the expressions of ACTB, C3, CXCL3, NQO1, and SELP genes between groups. ACTB expression in LET2 was higher than that in LET1 (Figure 7A, *P* < 0.05). The expressions of C3 (Figure 7B) and CXCL3 (Figure 7C) in LET2 were higher than that in Con2 (*P* < 0.05). The expression of NQO1 in LET1 was higher than that in Con1 and LET2 (Figure 7D, *P* < 0.05). The expression of SELP in LET1 was higher than that in Con1 (Figure 7E, *P* < 0.05).

**Figure 7.**
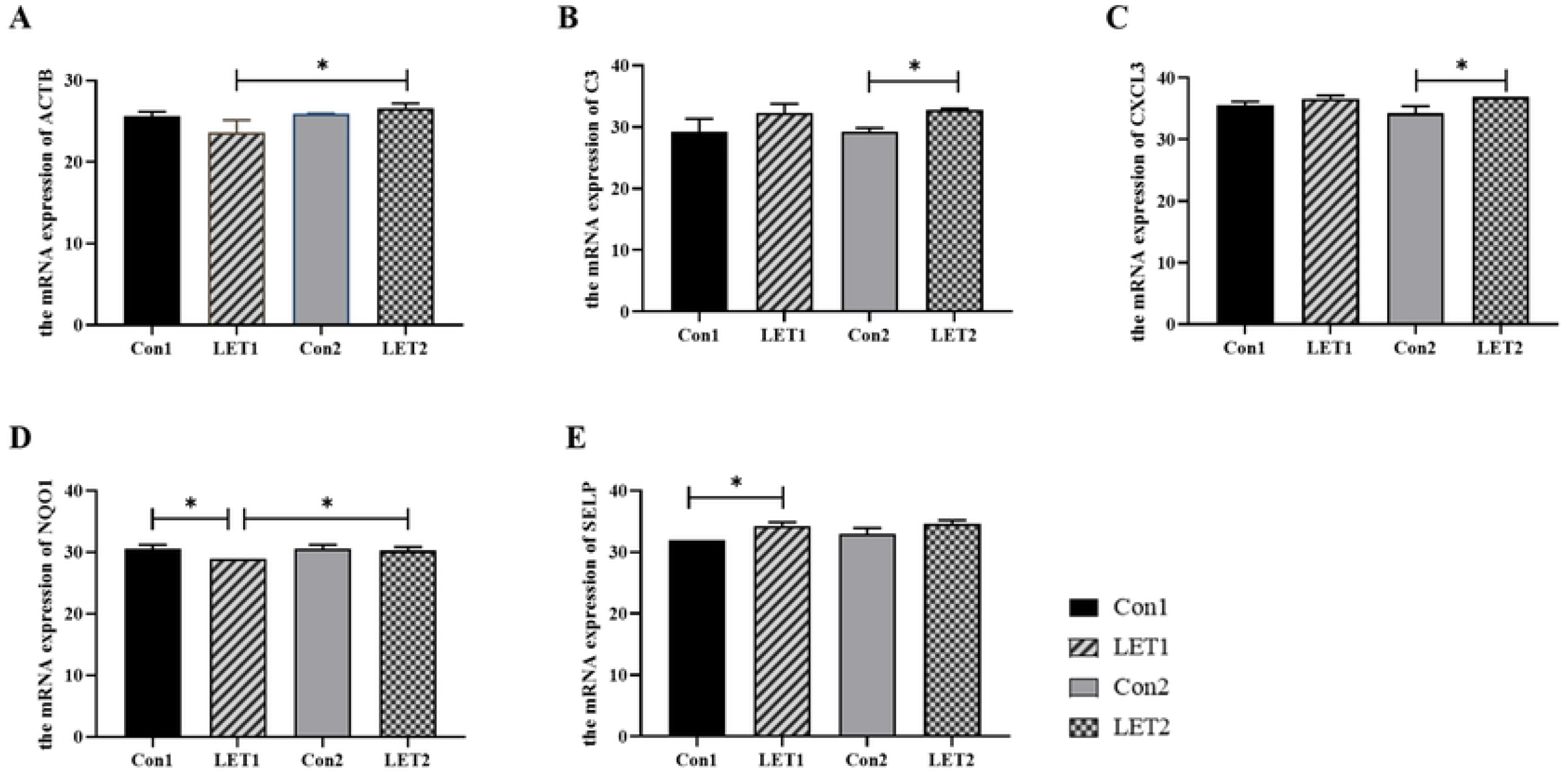

## 4 Discussion

PCOS is a complex reproductive and endocrinal disease which the pathogenesis remains unclear. The current treatments are mainly focused on symptomatic approaches rather than curative. Importantly, exploring the pathological mechanisms of PCOS is an urgent to be addressed. Our previous study indicates that the continuous administration of LET, 200μg/d, to female rats for 12 weeks before puberty results in a PCOS model with both the reproductive and metabolic phenotypes [18]. This modeling method has also been used in many PCOS researches [29–31]. Recent studies indicated that increased cytokines were observed promotes in PCOS patients [32,33]. In order to further understand the association between pathological process of PCOS and inflammation, the changes of inflammatory response and metabolic characteristics in PCOS rats were investigate at 6 and 12 weeks after the LET intervention in this study. It was found that the state of high inflammatory status occurred before PCOS was developed, and this inflammation was also accompanied with obesity, accumulation of inguinal fats, hyperglycemia, and abnormal ovarian morphology. Therefore, our study indicated that a proinflammatory state might be an important factor in contributing to the development of PCOS.

TLR4 belongs to the Toll-like receptor family and expresses in a variety of cells in the body. TLR4 can initiate innate immune responses, modulate adaptive immunity, and defense against pathogens through recognition of pathogen-associated molecular patterns [34]. NF-κB, an important nuclear transcription factor, is a key downstream signal molecule in TLR4 signaling pathway [35], and can initiate the transcription of genes involved in the inflammation and immune responses [36]. In order to investigate the changes of inflammation during the pathogenesis of PCOS, this study examined TLR4 and NF-κB in ovaries and inguinal fat at different timepoints in LET-induced PCOS rats. It is known that NF-κB can bind to inhibitor of κB (IκBα) in the cytoplasm, but once dissociated from IκBα [37,38], allowing NF-κB to enter the nucleus, and NF-κB binds to target genes’ promoter regions, thereby triggering the expression of many genes involved in immune and inflammatory responses [39].Therefore, in the present experiment, we examined the expression levels of NF-κB in the nucleus and cytoplasm in inguinal fats and ovarian tissues, respectively. The results revealed that high expression levels of TLR4 and NF-κB in cytoplasm were found in both inguinal fats and ovarian tissues of rats at either 6 or 12 weeks of LET intervention, and the expression levels were higher at 12 weeks than at 6 weeks. However, NF-κB in nucleus expression was low in inguinal fats and ovarian tissues at 6 and 12 weeks of LET intervention and was even lower at 12 weeks than at 6 weeks. Differentially expressed levels of NF-κB in nucleus and cytoplasm level may represent the activation of TLR4/NF-κB signaling pathway and indicate the occurrence of inflammatory responses in inguinal fats and ovarian tissues. At present, more studies have confirmed that some drugs, such as Shaoyao-Gancao Decoction [40], Astragaloside IV [41] and Soy isoflavones [42], could reduce the inflammatory response in PCOS rats through modulating the NF-κB/TLR4 pathway; the effect is associated with altering gut microbiome structure, regulating secretion of proinflammatory cytokines and enhancing anti-inflammatory capacity. Our results also provide a theoretical basis for future drugs to treat PCOS by modulating the mechanism of TLR4/NF-κB in inguinal fats and ovarian tissues.

In PCOS patients, obesity is a common characteristic with an estimated prevalence of 49% and the risk is 2.8-fold higher among women with PCOS than healthy women [43]. As obesity is often linked to chronic low-grade inflammation, inflammation may be a contributing factor to obesity-related complications [44]. In this study, we found that the body weight of the PCOS rats was higher than that of the control group from the 5^th^ weeks of LET intervention until 12^th^ weeks. However, since this experiment only measured the inflammation level in the PCOS rats at 6 and 12 weeks of LET intervention, the relationship between body weight and inflammation changes during the development of PCOS could not be fully determined. But some studies have shown that inflammation was not only associated with obesity, but increased levels of neonatal systemic inflammation was reported to precede the onset of obesity [45–47]. Moreover, in the treatment of obesity, anti-inflammation has been identified as an important strategy in obesity-related metabolic syndromes [48,49]. Therefore, the interactions between obesity and low-grade inflammation during PCOS need to be further studied.

Oh *et al*. proposed that women with PCOS showed a ‘male-like’ fat distributions with excessive abdominal (inguinal and visceral) fat accumulation [50]. Adipose tissue is the main source of producing cytokines [51,52]. There was growing evidence indicating obese mice on high fat diets had higher levels of inflammatory cytokines (e.g., TNF-a, interleukin-1β, IL-6) in their livers and adipose tissues [53,54]. Furthermore, hyperglycemia itself is a mediator of inflammation [55], which can also contribute to increasing inflammatory markers including TNF-α and IL-6 [56]. NF-κB can also be activated and stimulated in response to cellular stress including hyperglycemia, obesity and oxidative stress [57]. Recent studies have shown that in obese PCOS subjects, the NF-κB expression was higher than in lean healthy subjects [58]. Also, obesity, especially the abdominal obese phenotype, plays a significant role in activating the NF-κB pathway [59]. In this study, we found both higher inguinal fat and blood glucose in PCOS rats at 6 and 12 weeks of LET intervention. According to another study, excessive inguinal fat was associated with impaired organ function and chronic inflammation in obese individuals [60]. This is also consistent with the high inflammatory response we observed in the inguinal fats of PCOS rats at 6 and 12 weeks of LET intervention. Furthermore, Lionett *et al*. identified that women with PCOS had lower levels of oxidation and mitochondrial respiration in abdominal and gluteal inguinal fats than healthy women [61]. However, more studies are needed to unravel the specific pathological changes in different adipose tissues among PCOS.

Although the etiology of PCOS remains unknown, mounting evidence suggests that follicular dysfunction is involved in causing the infertility in women with PCOS [62]. PCOS patients have increased cystic follicles, a thickened thecal cell layer, loose arrangement of granular cells, and reduced corpus luteum; animal models of PCOS also showed similar alterations [62,63]. In this study, follicles with numerous follicular cysts were only present in the ovaries of LET2 group compared with other 3 groups. In addition, the LET2 group showed no fluctuation in the body temperature of PCOS rats during 7-8 and 10-12 weeks of LET intervention, which is also an indication of poor ovulation in PCOS rats. Our study indicates that it may be related to the presence of an inflammatory microenvironment in the ovaries. In the ovary, cytokines are secreted by leukocytes, oocytes and follicular cells [64]. These cytokines are involved in multiple biology processes including regulating the synthesis of gonadal steroids folliculogenesis, ovarian cells proliferation, and the function of corpus luteum [65,66]. The increased inflammation in PCOS ovaries has also been reported to impact follicular growth, and TLR4/NF-κB activation creates an inflammatory environment in the ovary that disrupts ovulation [67]. Interestingly, the development of follicles in the LET1 group showed no different compared with control groups, although there were high expression levels of TLR4 and NF-κB, it may be due to the inflammation intensity and exposure time are not strong and long enough to induce an adverse effect on ovaries, which needs further research.

NF-κB plays a critical role in the initiation and resolution of inflammation [68] and can been activated and translocate into the nucleus to upregulate transcription of several inflammatory genes [69,70]. We showed, for the first time, that five differentially expressed NF-κB signaling target genes in ovarian tissues in LET-induced PCOS rat at two different timepoints, specifically, ACTB, C3, CXCL3, NQO1 and SELP. So far, only a few studies have explored their pathologic mechanisms in PCOS. Huang team performed cDNA microarray and qRT-PCR analysis of cumulus cells isolated from PCOS patients, and their results suggested that CXCL3 was related to oocyte nuclear maturation in PCOS patients [71,72]. In order to determine the candidate genes of PCOS, Shen et al. performed comprehensive analysis using GSE345269 microarray date which consisted of 7 granulosa cell samples of PCOS patients and 3 healthy normal granulosa cell samples [73]; it was found that ACTB were upregulated in PCOS patients but the difference was not statistically significant. The complement factor C3 is a pivotal component of inflammation; studies found an association of serum C3 and Creactive protein (CRP) with insulin resistance in PCOS subjects [74,75]. Studies also identified that NQO1 protein expression was significantly increased in the endometrium of women with PCOS and this change was similar to those found in patients with endometrial cancer [76]. Further functional experiments will be needed to investigate the role of these genes in PCOS, which is also the main focus of our future experiments.

There are some limitations in this study. Samples can be collected and observed at more time points during LET intervention, such as 4 weeks and 8 weeks. On the other hand, we could analyze the inflammatory status in different tissues, such as liver, visceral fats and hypothalamus, and more comprehensively observations of the inflammatory processes in PCOS are needed to have a better understanding of how inflammation contributes to the development of PCOS.

## 5 Conclusions

This study, for the first time, investigate the associations of inflammation status and PCOS characteristics across two different timepoints. Activation of the TLR4/NF-κB pathway in inguinal fats and ovarian tissues persisted during the formation of LET-induced PCOS in rats. Moreover, ACTB, C3, CXCL3, NQO1 and SELP may be key regulatory target genes in ovarian tissue. Also, the high inflammatory state in PCOS may induce or exacerbate the phenotypic features of PCOS such as hyperglycemia, weight gain, adipose tissue accumulation and follicular dysplasia. Therefore, improving inflammation may play a critical role in preventing or attenuating PCOS. The regulation of the TLR4/NF-κB pathway could be new target in the prevention and treatment of PCOS.

## Notes

### Competing Interest Statement

The authors have declared no competing interest.

